# LRP2: A proteogenomics pipeline for long-read informed protein isoform analysis and discovery

**DOI:** 10.64898/2026.05.27.728216

**Authors:** Megan D. Schertzer, Julia T. Lewandowski, Emily F. Watts, Will Rosenow, Madison M. Mehlferber, Erin D. Jeffery, Scott I. Adamson, Jocelyne Bruand, Elizabeth Tseng, Yaseswini Neelamraju, Francine E. Garrett-Bakelman, Egor Dolzhenko, David A. Knowles, Gloria Sheynkman

**Author notes:** Corresponding authors. Megan Schertzer, David Knowles, Gloria Sheynkman. Equal contribution (Megan D. Schertzer, Julia T. Lewandowski).

## Abstract

Most human genes produce multiple RNA isoforms, yet it remains unclear which isoforms are translated into stable, functional proteins. Long-read RNA sequencing resolves full-length transcript structures and, when paired with mass spectrometry, can provide empirical evidence of isoform translation. Despite this opportunity, comprehensive workflows integrating isoform discovery, open reading frame prediction, peptide identification, and protein inference remain limited, leaving users to handle these steps piecemeal. Here, we present LRP2, a modular, end-to-end long-read proteogenomics pipeline built in Nextflow. LRP2 scales transcript discovery to hundreds of samples via PacBio’s latest Isocall tool, removes technical artifacts with SQANTI QC, generates and classifies predicted proteomes via CPAT and SQANTI Protein, performs multi-group differential expression and usage analysis via edgeR, DRIMSeq, and a long-read adaptation of LeafCutter, and integrates protein-level evidence from DDA and DIA MS data through FragPipe. For cross-dataset comparison of novel isoforms, LRP2 employs deterministic splice-junction, coordinate-based isoform identifiers. Used as an integrated pipeline, LRP2 enables the detection of novel peptides and improves the protein isoform inference to confirm protein isoform translation.

**Availability and implementation:** LRP2 is freely available as a modular Nextflow pipeline at https://github.com/sheynkman-lab/LRP2 (v2.0.0, archived at https://doi.org/10.5281/zenodo.22795865). LRP2 supports Docker, Apptainer, and Conda environments with GENCODE references.

**Contact:** Megan Schertzer

David Knowles

Gloria Sheynkman

## 1 Introduction

Alternative promoter usage, polyadenylation, and splicing generate extensive transcriptome complexity and expand the potential diversity of protein products (Nilsen and Graveley 2010; Kelemen et al. 2013). Alternative transcript isoforms are produced from nearly all multi-exon human genes, and their usage varies across cell types, developmental context, and disease states (Pan et al. 2008; E. T. Wang et al. 2008; Baralle and Giudice 2017; Scotti and Swanson 2015).

Advances in long-read sequencing (LRS) have enabled increasingly complete catalogs of full-length transcript isoforms across diverse cellular conditions (Pardo-Palacios, Wang, et al. 2024; Reese et al. 2023). However, transcript-level catalogs alone cannot determine which isoforms are translated and stably expressed as proteins, which are the ultimate effectors of many splicing-driven functional changes. Consequently, the extent to which transcript diversity contributes to protein diversity is still unclear.

Proteogenomics addresses this longstanding problem by integrating transcriptomic and proteomics measurements. In long-read proteogenomics workflows, LRS-derived transcriptomes are used to predict full-length protein sequences that can be compiled into sample-specific protein databases for mass spectrometry (MS) analysis (Nesvizhskii 2014; Sheynkman et al. 2016). This approach identifies isoform-informative peptides, including novel peptides absent from reference databases, to validate protein-level expression (Riepe et al. 2024).

While consortia are starting to generate matched LRS and MS data, there is a scarcity of scalable, end-to-end frameworks that integrate transcript discovery, open reading frame (ORF) prediction, and protein identification. Instead, analysis is conducted by manual integration of individual tools. Many tools are available for LRS analysis alone (Monzó et al. 2025), performing tasks ranging from quality control to gene annotation, though they remain largely focused on transcript-level analyses, not downstream protein-level outcomes. Similarly, ORF prediction tools can assess coding potential of novel transcript sequences, but typically operate as standalone modules, decoupled from downstream protein database generation. Finally, existing proteogenomic workflows construct customized protein databases by integrating transcript-derived sequences with MS data. However, these approaches are predominantly designed for short-read RNA-seq, limiting resolution to local splice events rather than full-length isoform structures (Cesnik et al. 2021; Zhu et al. 2025). Only a handful of recent tools support long-read proteogenomics, including our original LRP pipeline (Miller et al. 2022; Kulej et al. 2026; Kore et al. 2026), but none were designed for multi-sample, multi-condition studies.

We previously developed a “long-read proteogenomics” pipeline, LRP, one of the first open-source frameworks to integrate long-read transcriptome data with matched MS for sample-specific protein isoform identification (Miller et al. 2022; Mehlferber et al. 2022). Here, we present LRP2, a ground-up rebuild of this pipeline designed to enable large-scale proteogenomic analysis across multiple samples and conditions. LRP2 is a modular Nextflow pipeline, deployable via Docker, Singularity, or Conda, and supports GENCODE annotations across human and mouse. To enable cross-study comparison of novel isoforms, LRP2 assigns hash-based identifiers based on each transcript’s splice junction chain, providing reproducible coordinate-based IDs independent of any single tool’s naming conventions.

## 2 Materials and methods

### 2.1 Nextflow pipeline architecture and design

LRP2 is a comprehensive redesign of our original pipeline architecture (Miller et al. 2022), prioritizing scalability, flexibility, and ease of use. The pipeline is organized into five modular subworkflows (S1-S5): transcript discovery, transcriptome characterization, proteome prediction, multi-condition differential analysis, and proteomics search (Figure 1A). The pipeline is containerized via Apptainer/Singularity for deployment on institutional HPC clusters, broadening accessibility to the academic community, while also maintaining Docker compatibility for usage in cloud computing environments. LRP2 has flexible data requirements: users may run with only LRS data (S1-S4), only MS data (S5), or paired LRS and MS samples (S1 - S5). Execution can begin at multiple entry points (S1, S2, S4, or S5), so analyses can be repeated or extended from existing outputs without regenerating upstream results. For example, users may rerun differential analysis with alternative sample groupings or parameters, or rerun a proteomics search with new MS fractionations against a proteome built in a previous run. Beyond the raw LRS and MS data themselves, the only required input is a CSV sample sheet specifying sample names, file paths, data type (RNA or protein), condition, and MS acquisition mode (DDA or DIA). Default parameters and configuration files are provided but may be customized by the user.

**Figure 1.**
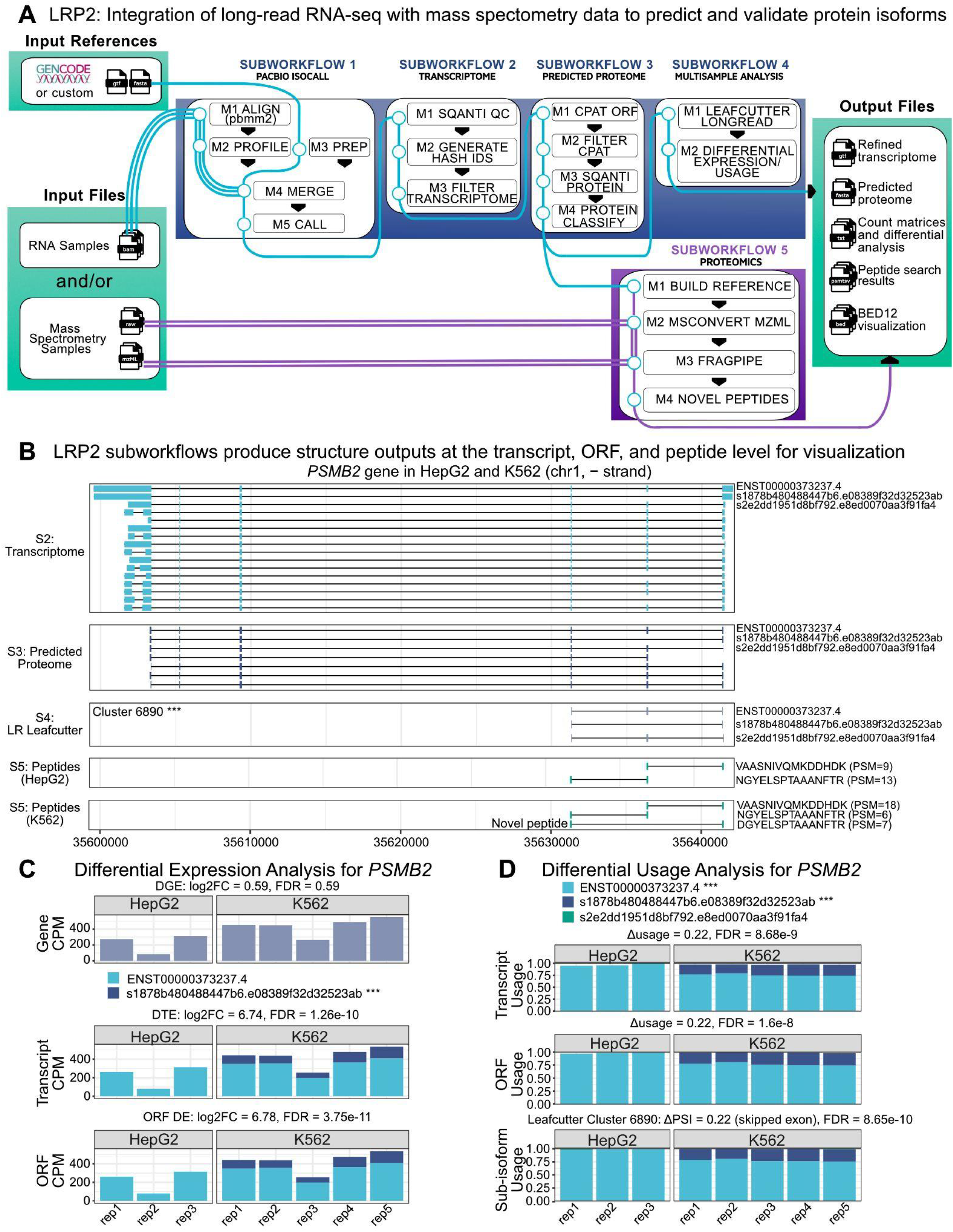
Overview of the LRP2 workflow. (A) Nextflow pipeline showing inputs, five modular subworkflows, and outputs. (B) All LRP2 subworkflows output GTF and/or BED12 files for coordinate-based visualization. Structure plots for an example gene, *PSMB2*, from the transcriptome, predicted proteome, multisample analysis, and proteomics subworkflows. For simplicity, only the three most abundant isoforms are labeled on the right– the ENST ID for the annotated isoform and hash IDs for novel isoforms. For the LeafCutter panel, the three sub-isoform structures of the significant cluster are plotted, and the peptide panels show the corresponding peptides detected in HepG2 and K562. Values in parentheses to the right of S5 are the number of peptide-spectrum matches (PSMs) supporting each peptide. K562 contains an additional novel peptide not detected in HepG2. (C) Differential expression analysis (edgeR) at the gene, transcript, and ORF level and (D) differential usage analysis (DRIMSeq and LR LeafCutter) at the transcript, ORF, and sub-isoform levels of *PSMB2* between HepG2 (n = 3) and K562 (n = 5) RNA biological replicates. After edgeR and DRIMSeq filtering, only two *PSMB2* isoforms are retained for testing. Transcript and ORF usage are calculated across all isoforms, but only the two retained by DRIMSeq are plotted; bars therefore do not sum to 1. After LeafCutter filtering, only three sub-isoforms from cluster 6890 are retained for testing. The cluster represents an exon-skipping event that overlaps an alternative 5’ splice site, though the exon-skipping event drives the changes seen at the isoform-level. Significance: *** FDR < 0.001.

### 2.2 Scalable transcript discovery and MS proteomics search

The original LRP pipeline was only suitable for single-sample analysis (Miller et al. 2022). LRP2 introduces parallelized execution across samples for read alignment and all proteomics modules. In addition, PacBio’s Isocall ‘profile’ and ‘call’ modules replace the ‘cluster’ and ‘collapse’ steps used in earlier IsoSeq-based workflows (Gordon et al. 2015), substantially reducing computational requirements for transcript discovery (Dolzhenko et al. 2026). For users to assess scalability, we have included runtime, CPU usage, and peak memory tracing for all individual modules, across 10-124 ENCODE4 LRS samples (Reese et al. 2023) and 1-51 jointly processed MS fractions (Sinitcyn et al. 2023) (Supplemental Note N1; Supplemental Tables 1-2). Across all four RNA subworkflows (S1-S4), LRP2 processed 124 samples in approximately 9 hours of runtime (excluding queue time). Although the proteomics subworkflow (S5) runs in parallel per sample, we evaluated scalability by jointly processing fractionated MS samples. LRP2 S5 completed 51 raw MS files in an average of 3 hours and 40 minutes, with Fragpipe taking the longest at 3 hours and 29 minutes using 346.4 GB peak memory on average against a combined ENCODE4 LRS (124 samples) and GENCODE V49 reference. Together, these results demonstrate that the updated LRP2 enables cohort-scale proteogenomics analysis.

### 2.3 Deterministic isoform identifiers

Annotated transcript isoforms have a stable accession (e.g., GENCODE ENST) that serves as a common key; novel isoforms have none. Cross-study comparison of novel transcript isoforms is hindered by the random, run-specific identifiers that most long-read assembly tools assign. To address this gap, LRP2 assigns each isoform identifier by hashing its internal splice junction coordinates with SHA-256 (GENE_NAME::junction_hash). Because the hash is deterministic, isoforms with identical splice-junction structures receive the same identifier across independent analyses. Transcription start and end coordinates are not included in the hash, because read starts and ends are subject to technical variation, and including these would break the reproducibility of the hash. Hash IDs provide a reference-free key for novel isoforms and reduce reliance on downstream transcript-merging procedures for cross-study comparison.

### 2.4 Module for differential analysis of genes, isoforms, and splicing events

LRP2 includes a new multisample analysis subworkflow that performs all pairwise comparisons among two or more conditions (e.g. wild type vs. knockout, normal vs. disease). Differential gene, transcript, and ORF expression–the latter aggregating transcripts with the same predicted coding sequence– are quantified using edgeR (Robinson et al. 2010), and differential transcript and ORF usage are identified using DRIMSeq (Nowicka and Robinson 2016).

In addition to transcript-level analyses, LRP2 performs differential splicing analysis using a long-read adaptation of LeafCutter (Li et al. 2017). Long-read LeafCutter is under active development and is included in LRP2 as a preliminary implementation (Supplemental Note N2). LeafCutter groups overlapping introns into clusters and tests for shifts in their relative usage between conditions. Our long-read adaptation preserves this intron clustering strategy and LeafCutter’s Dirichlet-multinomial model, while extending it to leverage information unique to LRS. Because long reads capture full transcript structures, the method resolves which junctions co-occur on the same molecule. This allows us to model *“sub-isoforms”*, i.e. the possible chains of exons and introns used within an intron cluster. Sub-isoforms provide an intermediate resolution between individual junctions and full transcripts, and are used instead of junctions as the unit of differential usage within each cluster.

Together, these modules test for differential expression and usage across a hierarchy of units–genes, transcripts, ORFs, and splice events (captured by sub-isoforms)–and run by default for each unique pair of conditions with two or more RNA samples, supporting automated pairwise comparisons at scale.

### 2.5 Integrated MS-based proteomics search

LRP2 integrates long-read-derived, sample-specific predicted proteomes directly into downstream MS search workflows. When run as part of the full pipeline, the proteomics module uses protein databases generated from matched or related long-read transcriptome samples. Alternatively, the proteomics workflow can be run independently using user-supplied protein FASTA databases. Proteomics searches are implemented through FragPipe-based workflows supporting both data-dependent (DDA) and data-independent acquisition (DIA) MS datasets (Kong et al. 2017; Yu et al. 2021, 2023). Default configurations are provided, but users are expected to adapt workflows and search parameters to their specific MS instrumentation and experimental setup. Users can specify complex proteomics experimental designs through the sample sheet, including fractionated samples, biological replicates, and repeated MS injections from the same biological sample.

### 2.6 Standardized outputs and genome browser compatibility

Each LRP2 subworkflow produces standardized outputs, including GTF, FASTA, BED12, and count matrix files, enabling downstream transcriptomic and proteomic analyses. The BED12 outputs support direct visualization of transcript, ORF, splice-junction, and peptide structures in genome browsers such as IGV and the UCSC Genome Browser.

### 2.7 Demonstration datasets

To demonstrate LRP2 in the Results section, we processed publicly available K562 and HepG2 LRS and MS data (Supplemental Table 1). Raw LRS FASTQ files for HepG2 (n = 3) and K562 (n = 5) were downloaded from ENCODE4 (Reese et al. 2023). Raw MS files from trypsin-digested, higher energy collisional dissociated (HCD), DDA runs of HepG2 (n = 51 fractions) and K562 (n = 34 fractions) were downloaded from ProteomeXchange (PXD024364) (Sinitcyn et al. 2023).

## 3 Results

### 3.1 End-to-end LRP2 run on K562 and HepG2 LRS and MS data

LRP2 generates a refined transcriptome, predicted proteome, and matched proteomics evidence from LRS and MS inputs (Figure 1A). The first three subworkflows (PacBio Isocall, Transcriptome, and Predicted Proteome) each generate a GTF of transcript structures, DNA or ORF FASTA, a BED12 file, and a count matrix. The fourth subworkflow (Multisample Analysis) produces LR LeafCutter and standard edgeR outputs alongside summary tables for DRIMSeq results across comparisons. The fifth subworkflow (Proteomics) generates standard FragPipe output files and a peptide summary table that maps peptides to isoforms and flags novel peptides. As a demonstration, we ran LRP2 on HepG2 and K562 LRS samples from ENCODE4 (Reese et al. 2023) and cell-line-matched MS data (Sinitcyn et al. 2023) using GENCODE v49 as a reference. Below, we present summary results across subworkflows and follow the gene *PSMB2* as a representative example throughout.

### 3.2 Transcriptome (S1-S2)

Transcript identification with Isocall produced 112,520 distinct transcript structures (i.e., splice junction chains) across 17,597 genes. Following SQANTI QC classification (Pardo-Palacios, Arzalluz-Luque, et al. 2024) and custom filtering, 93,687 transcripts (83.3%) were retained (Supplemental Figure S1A). Custom filters removed transcripts from non-protein-coding genes (n = 10,011), potential internal priming artifacts (n = 7,432), potential template-switching artifacts (n = 1,277), and transcripts from atypical SQANTI structural categories (Genic and Fusion; n = 117) (Supplemental Figure S1B). The transcriptome subworkflow retained 16 *PSMB2* transcripts with distinct splice junction chains. The canonical isoform (ENST00000373237.4) was the most abundant across all samples (Figure 1B, Transcriptome).

### 3.3 Predicted proteome (S3)

CPAT (L. Wang et al. 2013) identified 432,712 ORFs across 93,687 transcripts (up to five ORFs per transcript, the CPAT default). Applying CPAT’s recommended human coding probability cutoff (>0.364), 121,165 ORFs were retained as probable protein-coding (Supplemental Figure S1C). Because multiple candidate ORFs may be identified per transcript, a single best-supported ORF per transcript was then selected based on a combination of criteria, including agreement with GENCODE annotations, higher coding probability, and a low number of upstream ATGs (n = 87,409; Supplemental Figure S1D). Additional details and evaluation of ORF calling and selection are in Supplemental Note N3. The best-supported ORF per transcript was used for all downstream analysis, including building the custom proteome FASTA databases for MS searches. SQANTI Protein and the custom filtering module (Miller et al. 2022) further curated candidate ORFs by removing nonsense-mediated decay (NMD) transcripts and predicted 5’-truncated transcripts, then collapsed transcripts that produce the same ORF into a final set of high-confidence ORFs. *PSMB2* isoform complexity was reduced to 7 distinct ORFs after removal of 9 predicted NMD transcripts (Figure 1B, Predicted Proteome).

### 3.4 Multisample Analysis (S4)

Multisample differential analysis between HepG2 and K562 identified widespread changes in expression and usage across genes, transcript isoforms, and ORFs, as well as in local splicing patterns. Differential expression analysis with edgeR (Robinson et al. 2010) identified 2,276 genes, 2,981 transcripts, and 2,482 ORFs as differentially expressed between HepG2 and K562 (FDR < 0.05; Supplemental Figure S1E, S1F). Differential transcript and ORF usage analysis with DRIMSeq (Nowicka and Robinson 2016) identified 576 transcripts and 384 ORFs with significant usage changes (FDR < 0.05; Supplemental Figure S1G). Long-read LeafCutter identified 747 differentially spliced sub-isoform clusters (FDR < 0.05; Supplemental Figure S1H).

*PSMB2* illustrates how these complementary differential analyses resolve isoform-level regulation not apparent from gene-level expression alone. Although total gene expression did not differ significantly between HepG2 and K562 (FDR = 0.59), a novel isoform was enriched in K562 relative to HepG2. This isoform was significant across transcript, ORF, and splicing-level differential analyses (FDR < 0.001; Figure 1C-D). Consistent with this, LR LeafCutter identified an exon-skipping event overlapping an alternative 5’ splice site (cluster 6890), indicating that this local splicing change underlies the observed isoform shift (Figure 1B, LR LeafCutter).

### 3.5 Proteomics (S5)

MS searches against the custom long-read-derived database using FragPipe’s standard DDA workflow detected 212,511 and 204,050 peptides in HepG2 and K562, respectively (Supplemental Figure S1I). Of these, 723 and 464 peptides in HepG2 and K562 were classified as novel, mapping exclusively to novel ORFs in the LRP2 custom database. For *PSMB2*, we identified a novel peptide unique to the exon-exclusion isoform, detected only in K562 at both the RNA and peptide levels (Figure 1B; Peptides; Supplemental Figure S2), supporting translation of the isoform.

## 4 Future Development

Several aspects of LRP2 are the focus of ongoing development. First, the Isocall Subworkflow (S1) is restricted to PacBio reads. We plan to add an alternative Oxford Nanopore (ONT)-compatible subworkflow, with comparable scalability as the chief design consideration. In the meantime, users with ONT data can pre-process raw reads using their own pipeline and input a GTF and count matrix into LRP2 at the start of the Transcriptome Subworkflow (S2), bypassing S1. Second, transcript identification is purely coordinate-based, so gene fusions, SNPs, and indels are not incorporated into the transcriptome, predicted proteome, or proteomics subworkflows. Third, the predicted proteome subworkflow selects a single best-supported ORF per transcript for FragPipe searches, which may miss alternative ORF products (Mudge et al. 2022). Beyond addressing these limitations, we will continue to expand functionality in the proteomics subworkflow. We will add support for TMT-based quantification in multiplexed experiments. We are also exploring perplexity-based metrics to improve protein inference by identifying the most parsimonious set of isoforms supported by peptide evidence (Schertzer et al. 2026). LRP2 has been developed following nf-core standards and will be submitted to the nf-core community pipeline repository to broaden LRP2’s utility for cohort-scale proteogenomic discovery.

## Acknowledgements

We thank everyone who contributed to the original LRP pipeline (Miller et al. 2022), which laid the foundation for this second version. We thank members of the Sheynkman Lab and the Knowles Lab for valuable feedback and testing during the development of LRP2. We gratefully acknowledge the University of Virginia Research Computing team for providing access to the Rivanna High-Performance Computing system and for their responsive support throughout this work.

