## Supplemental Figures for "LRP2: A proteogenomics pipeline for long-read informed protein isoform analysis and discovery"

**Supplementary Information**

### Supplemental Tables

**Supplemental Table S1: Data Overview.** Metadata and URLs for all 124 ENCODE4 LRS samples and ProteomeXchange HCD DDA MS data for 34 K562 and 51 HepG2 fractionations.

**Supplemental Table S2: LRP2 ENCODE4 Resource Summary.** Per-task Nextflow trace metrics for LRP2 execution on 124 ENCODE4 LRS samples and 51 HepG2 fractionated MS samples. For all modules in S1 - S4 (PACBIO_ISOCALL, TRANSCRIPTOME, PREDICTED PROTEOME, MULTISAMPLE_ANALYSIS), mean measurements were computed over 5 replicates, and for S5 PROTEOMICS, mean measurements were computed over 3 replicates. Included columns are task_id (Nextflow task identifier), name (concatenated task and sample filename), exit (exit code of process), submit (task submission timestamp), mean_duration (average total process time including queue waittime), mean_realtime (average actual process execution time), mean_%cpu (average CPU utilization), mean_peak_rss (average peak resident set size), mean_peak_vmem (mean peak virtual memory), mean_rchar (average bytes read), and mean_wchar (average bytes written).

### Supplemental Notes

**Supplemental Note N1: Resource requirements and scalability of LRP2 by subworkflow and module.** Runtime, memory, and CPU utilization measured across subsets of 10–124 ENCODE4 LRS samples and 1–51 HepG2 MS fractions, with process tag definitions and guidance on adjusting resource allocations.

**Supplemental Note N2: Long-read adaptation of LeafCutter.** Definition of sub-isoforms and how clustering and differential usage testing compares to the original short-read implementation.

**Supplemental Note N3: Evaluation of CPAT ORF calling and LRP2 selection accuracy against GENCODE annotation.** Accuracy of CPAT ORF calling and of the LRP2 best-supported ORF criteria against MANE and non-MANE transcripts, and sensitivity of selection accuracy to the coding probability threshold.

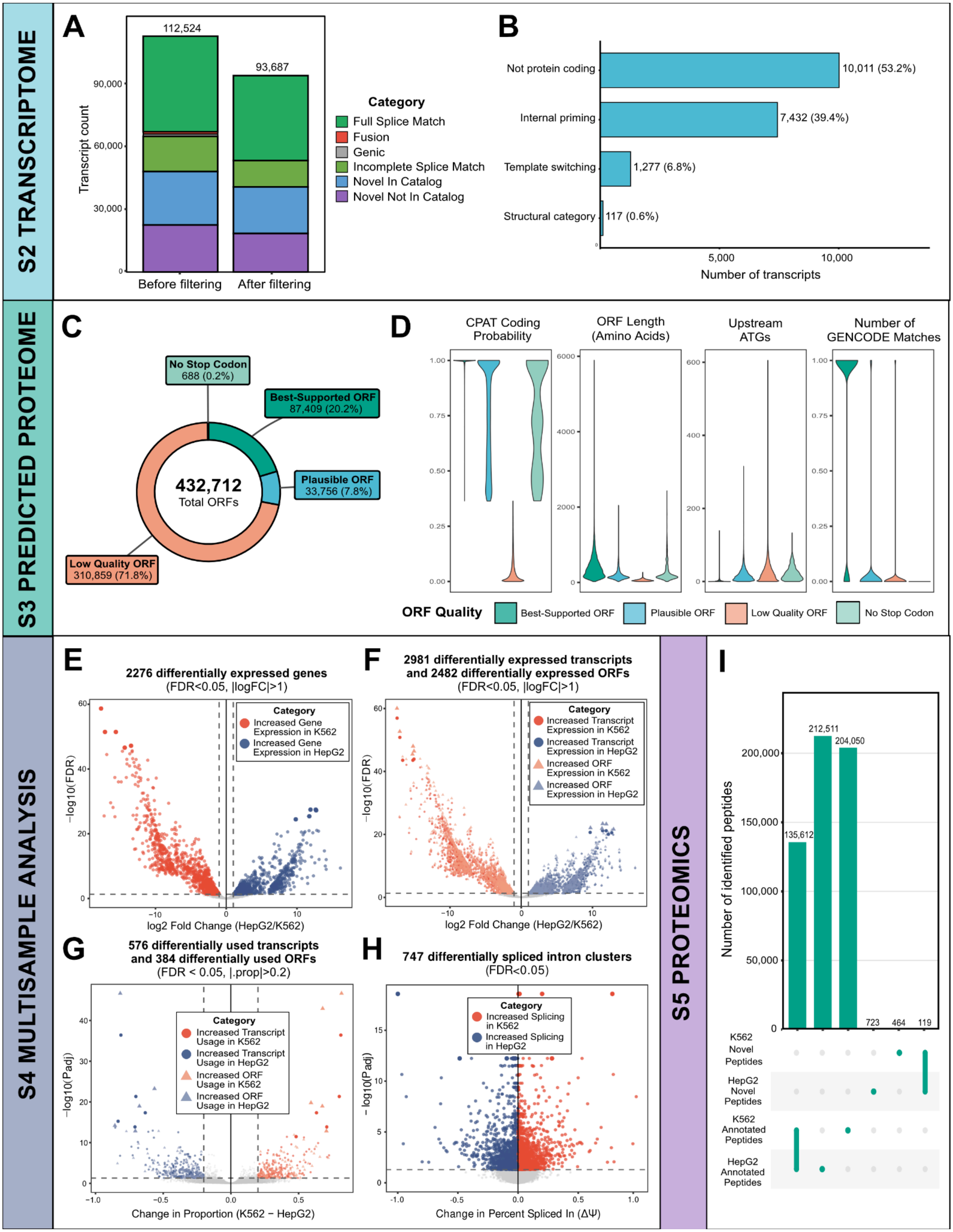


**Supplemental Figure S1. Summary of LRP2 output for K562 and HepG2 paired LRS and MS data.** (A) Transcript counts before and after transcriptome subworkflow filtering, categorized by SQANTI3 structural classification. (B) Breakdown of 18,841 filtered transcript removed by transcriptome subworkflow filtering, by category. (C) CPAT classification across 432,712 predicted ORFs. (D) ORF quality metrics stratified by predicted proteome subworkflow, stratified by predicted category; best-supported ORFs show higher coding probabilities, longer lengths, fewer upstream ATGs, and more GENCODE matches than lower-quality categories. (E) Volcano plot of differential gene expression; 2,275 genes were differentially expressed (FDR < 0.05, $|\log_2 FC| > 1$). (F) Volcano plot of differential transcript and ORF expression; 2,981 transcripts and 2,484 ORFs were differentially expressed (FDR< 0.05, $|\log_2 FC| > 1$). (G) Volcano plot of differential transcript and ORF usage; 405 transcripts and 265 ORFs were differentially used (FDR < 0.05, $|\Delta \text{proportion}| > 0.2$). (H) Volcano plot of differential splicing from Long-read LeafCutter; 2,845 intron clusters were differentially spliced (FDR < 0.05). (I) UpSet plot of unique peptides detected across cell lines. 212,511 and 204,050 annotated peptides were detected in K562 and HepG2, respectively (135,612 shared), with 723 and 464 novel peptides (119 shared).

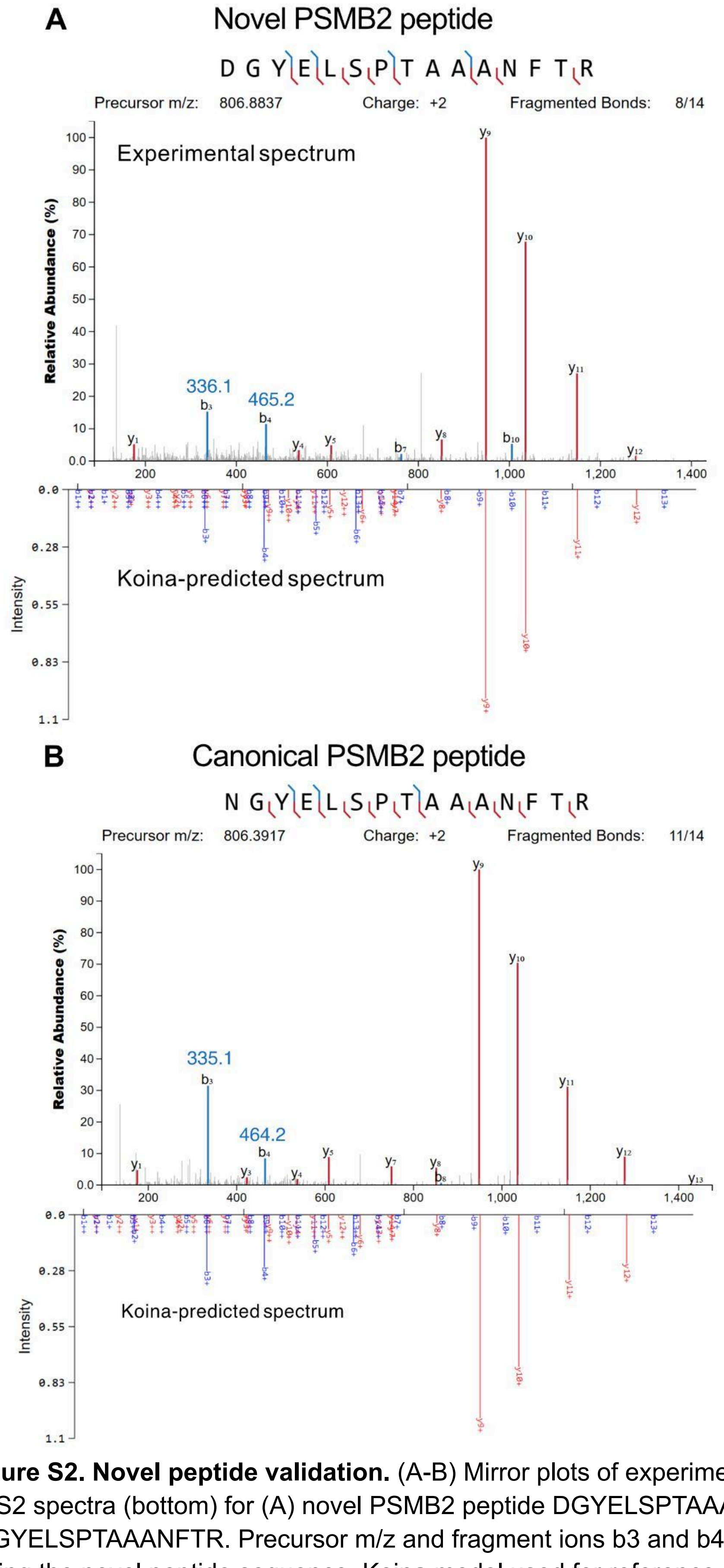


**Supplemental Figure S2. Novel peptide validation.** (A-B) Mirror plots of experimental MS2 spectra (top) and Koina-predicted MS2 spectra (bottom) for (A) novel PSMB2 peptide DGYELSPTAAANFTR and (B) canonical PSMB2 peptide NGYELSPTAAANFTR. Precursor m/z and fragment ions b3 and b4 are the distinguishing features for assigning the novel peptide sequence. Koina model used for reference spectra is Prosit_2020_intensity_HCD (Lautenbacher et al. 2025).
