## Supplemental Notes for "LRP2: A proteogenomics pipeline for long-read informed protein isoform analysis and discovery"

**Note N1: Resource requirements and scalability of LRP2 by subworkflow and module**

*Full Pipeline (S1-S5) Evaluation on 124 RNA Samples and 51 MS Fractions*

To benchmark LRP2 scalability for RNA Subworkflows S1-S4, we tested on subsets of the human ENCODE4 long-read RNA-sequencing datasets (Reese et al. 2023) of sample sizes N = [10, 20, 30, 40, 50, 65, 80, 95, 110, 124] (Table N1.1).

**Table N1.1.** Breakdown of cumulative number of reads and total dataset size in GB for each subset of the ENCODE4 LRS samples used for the S1-S4 scalability benchmark.

| Number of samples | Total Raw Reads | Total Dataset Size (GB) |
|---|---|---|
| 10 | 6,039,318 | 2.18 |
| 20 | 13,026,345 | 5.38 |
| 30 | 21,104,083 | 9.31 |
| 40 | 30,311,332 | 14.19 |
| 50 | 42,597,270 | 20.63 |
| 65 | 65,594,239 | 32.66 |
| 80 | 92,280,938 | 46.98 |
| 95 | 119,743,932 | 62.85 |
| 110 | 156,510,587 | 81.09 |
| 124 | 195,307,478 | 104.74 |

All measurements were performed on Intel Xeon E5-2690 v4 @ 2.60GHz nodes (56 cores, 220GB RAM) held under exclusive allocations. For each sample subset, 5 replicate measurements were taken in serial. Work directories were completely purged between replicates to additionally ensure full independence between runs. Runtime (minutes), CPU usage, and Peak Memory Usage (according to Nextflow execution trace `peak_rss`) are reported for each Module or Subworkflow in the following Figures.

In general, runtime for each subworkflow grows approximately linearly with N samples over the tested range, with no subworkflow showing evidence of super-linear scaling (Figure N1.1). In total, LRP2's S1-S4 RNA subworkflows completed analysis of the 124 samples in approximately 9 hours of runtime, excluding queue time, with the S3 subworkflow taking the longest.

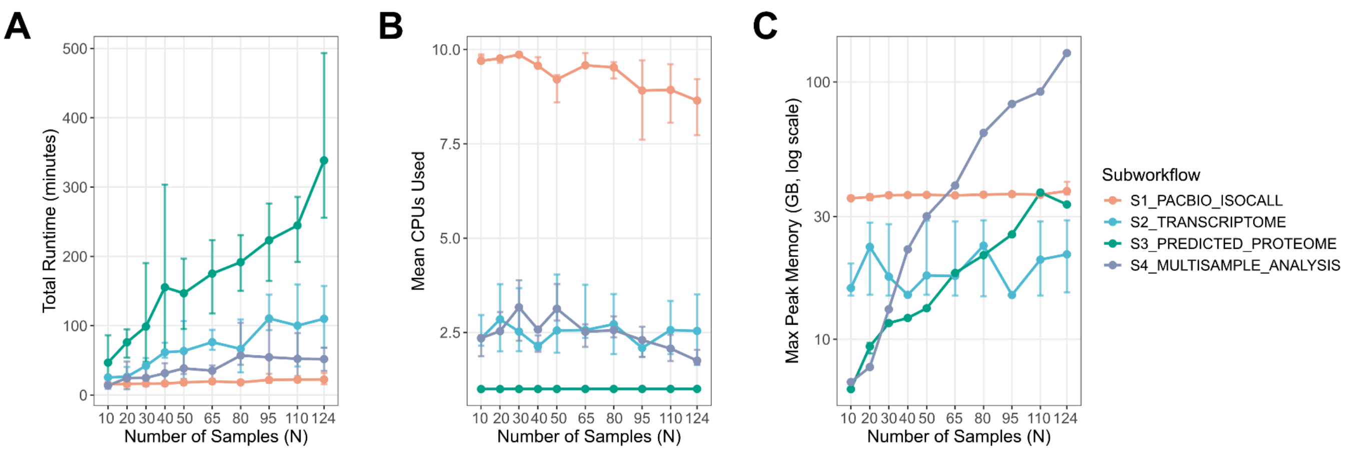


**Figure N1.1**. Overview of runtime and memory usage by LRP2 RNA subworkflows S1 - S4.

To evaluate LRP2 scalability for the proteomics subworkflow S5, we assessed runtime and resource usage as the number of fractionations (representing greater coverage) increased, because proteomics searches run in parallel for each unique mass spectrometry sample. We ran LRP2 from the S5 entry point, using a custom protein FASTA and CDS GTF input derived from a prior full pipeline execution, on eleven subsets of HepG2 trypsin-digested DDA mass spectrometry fractions (Sinitcyn et al. 2023) for N = [1, 5, 10, 15, 20, 25, 30, 35, 40, 45, 51] (Table N1.2).

**Table N1.2** Breakdown of total sample size in GB per number of MS fractionations used for each S5 scalability benchmark.

| Number of MS fractionations | Total Size (GB) |
|---|---|
| 1 | 1.78 |
| 5 | 7.87 |
| 10 | 15.44 |
| 15 | 22.87 |
| 20 | 30.44 |
| 25 | 37.68 |
| 30 | 44.68 |
| 35 | 51.71 |
| 40 | 58.49 |
| 45 | 65.14 |
| 51 | 73.92 |

All measurements were performed using identical hardware as the S1 - S4 benchmarking (Intel Xeon E5-2690 v4 @ 2.60GHz nodes, 56 cores, 220GB RAM) with each node held under an exclusive allocation, with 3 replicate measurements taken in serial for each fraction count. All modules run once per unique MS sample name in the samplesheet and are thus parallelized. The most significant memory bottleneck is proteomics search via M3_FRAGPIPE, and the FragPipe search time and memory requirements will scale relative to the number of fractionations per sample and complexity of the proteome (Figure N1.2).

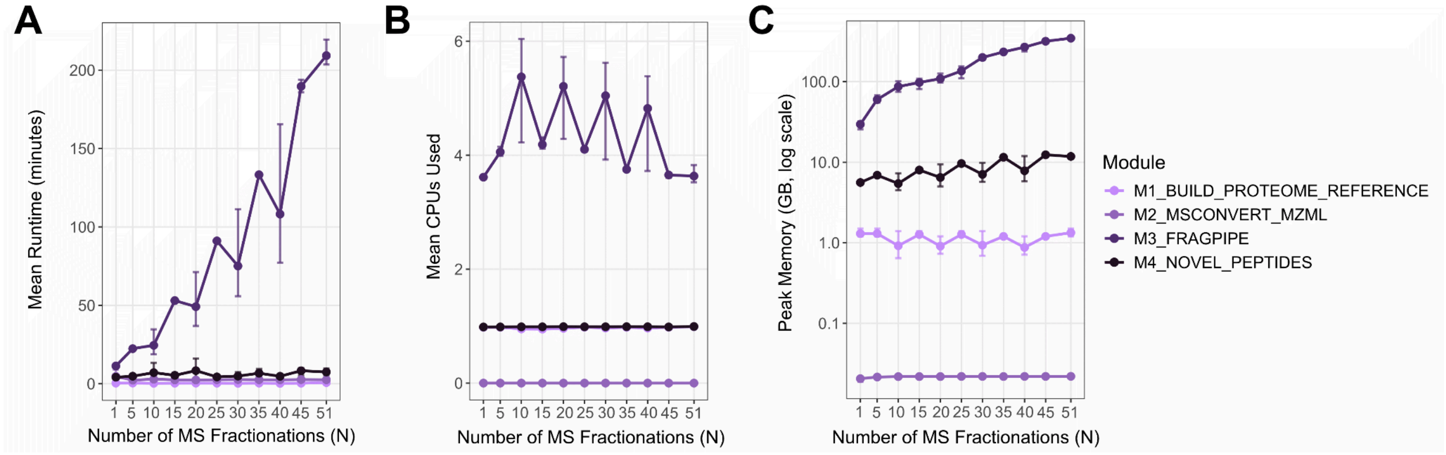


**Figure N1.2.** Overview of runtime and memory usage by LRP2 Proteomics Subworkflow S5.

*Preset Process Tags to Specify Module Resource Allocations*

In LRP2, each module within each subworkflow is associated with a process tag (Table N1.3) specified on line 3 of each module definition file (e.g. `modules/local/isocall_prep/main.nf`). We have assigned process tags to modules based on our empirical observations. Process tag definitions are found in `conf/base.config` for runs using Singularity/Apptainer, Docker, or Conda. Users analyzing larger datasets can modify the resources (`cpus`, `memory`, `time`) requested per process tag by directly editing that process tag's definition code block in this configuration file.

**Table N1.3.** LRP2 process tag labels found in conf/base.config.

| Nfcore process tag | CPUs | Memory (GB) | Time (hour) | Modules |
|---|---|---|---|---|
| process_single | 1 | 6 | 1 | S1_M1_ISOCALL_PREP<br>S1_M4_ISOCALL_MERGE<br>S5_M1_BUILD_REFERENCE |
| process_low | 1 | 16 | 1 | S1_M3_ISOCALL_PROFILE *<br>S1_M5_ISOCALL_CALL *<br>S2_M2_GENERATE_HASHIDS<br>S3_M1_CPAT_ORF |
| process_medium | 1 | 48 | 2 | S2_M3_FILTER_TRANSCRIPTOME<br>S3_M2_FILTER_CPAT<br>S3_M4_PROTEIN_CLASSIFICATION<br>S4_M2_DIFFERENTIAL_EXPRESSION<br>S5_M2_MSCONVERT_MZML<br>S5_M4_NOVEL_PEPTIDES |
| process_high | 16 | 128 | 3 | S1_M2_PBMM2_ALIGN<br>S2_M1_SQANTI_QC<br>S4_M1_LEAFCUTTER_LONGREAD |
| process_high_memory | 12 | 500 | 8 | S5_M3_FRAGPIPE |
| process_long | 1 | 32 | 8 | S3_M3_SQANTI_PROTEIN |

* Resource requests for these modules are overridden by module-specific `withName` directives in `conf/base.config`: ISOCALL_PROFILE uses 4 CPUs, and ISOCALL_CALL uses 12 CPUs, 5 GB, and 1 hour.

*Module-level Scalability Across 124 RNA Samples*

In S1, M2_ISOCALL_PROFILE and M3_ISOCALL_ALIGN run in parallel for each sample, so resources do not scale with the number of samples but rather the size of the individual samples (Figure N1.3). M3_ISOCALL_ALIGN uses PacBio's minimap2 aligner (`pbmm2`) and requires the most runtime, CPUs, and memory. If processing LRS samples with > 20 million raw reads, users may need to (a) increase memory allocation and/or (b) increase CPUs allocated to speed up alignment; as per pbmm2's documentation, it is recommended to provide more memory to fewer sorting threads than less memory to many sorting threads to avoid disk I/O contention.

M4_ISOCALL_MERGE and M5_ISOCALL_CALL run jointly across all samples, so resources scale with the number of samples. However, resource requirements are minimal; at 124 LRS samples, both modules complete in under 5 minutes with less than 1 GB peak memory.

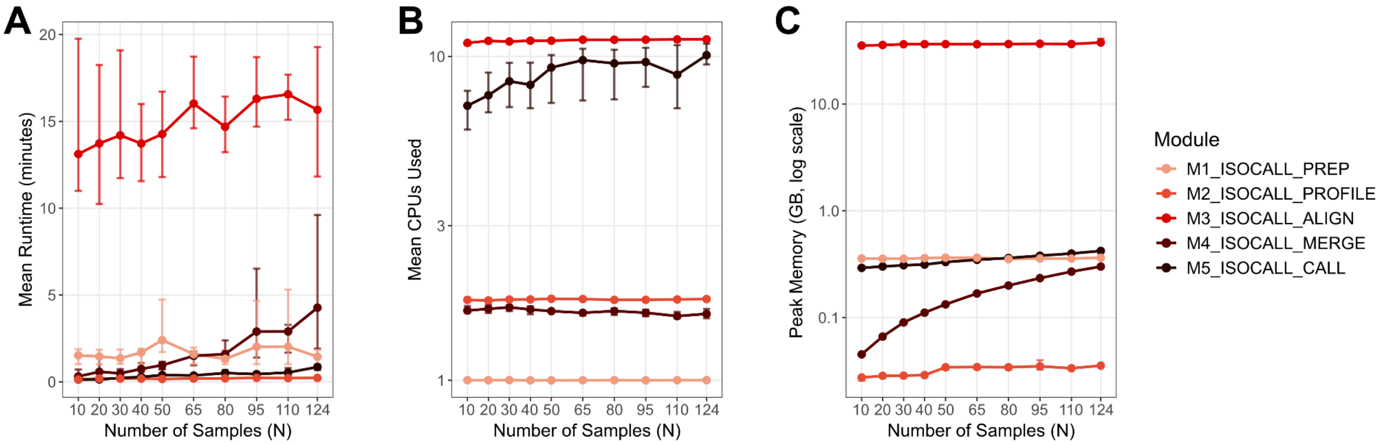


**Figure N1.3.** Subworkflow S1 PacBio Isocall.

In S2, all samples are jointly processed into a single refined transcriptome (Figure N1.4). M1_SQANTI_QC dominates runtime and resource usage, but does not scale linearly with N, which suggests that performance is more directly influenced by transcriptome complexity than sample count. We have implemented SQANTI QC with chunking, whereby the transcriptome is split into N chunks to be processed in parallel, where N number of CPUs allotted to the process. Therefore, for users analyzing large cohorts, it may be worth increasing the number of CPUs allocated to SQANTI QC. M2_GENERATE_HASHIDS and M3_FILTER_TRANSCRIPTOME, by contrast, demonstrate more modest, proportional increases in resource usage as sample size increases.

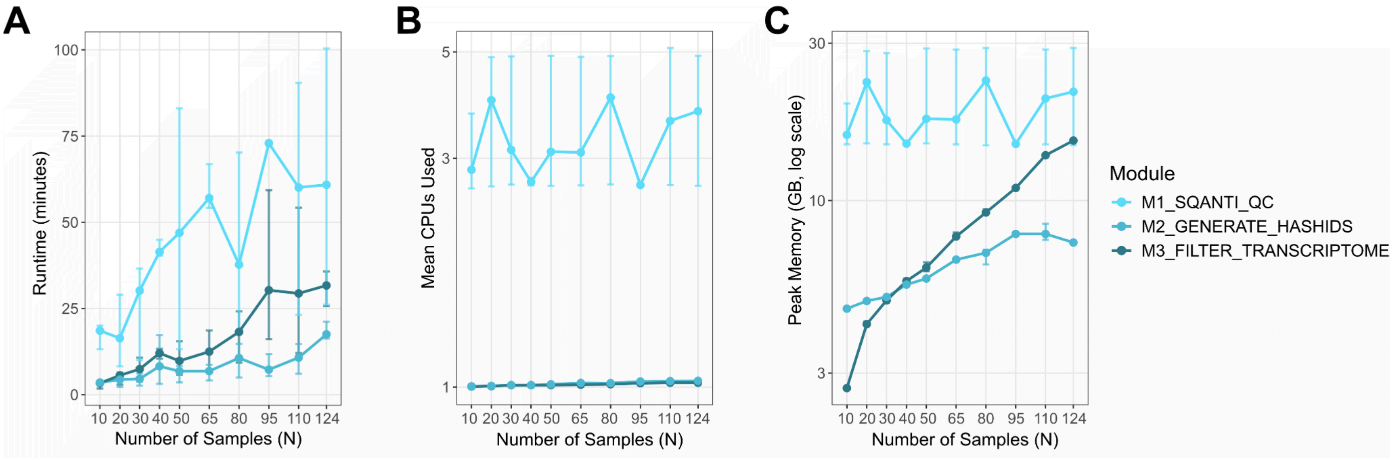


**Figure N1.4.** Subworkflow S2 Transcriptome.

In S3, the transcriptome output from S2 is processed as a single input summarizing all samples. We see that M3_SQANTI_PROTEIN and M4_PROTEIN_CLASSIFICATION modules require the most resources (Figure N1.5), and we intend to improve efficiency here as a future development. In the meantime, users may consider increasing max runtime allotted to `process_long` if processing complex transcriptomes.

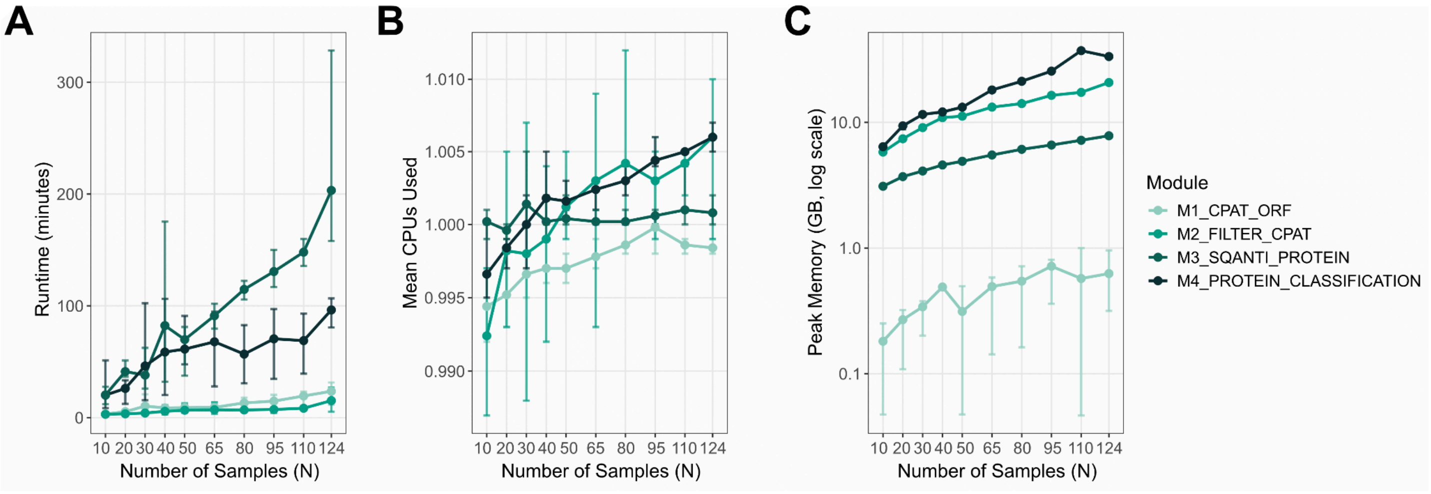


**Figure N1.5.** Subworkflow S3 Predicted Proteome.

In S4, M1_LEAFCUTTER_LONGREAD benefits from increased memory as sample sizes increase, while runtime scales modestly. M2_DIFFERENTIAL_EXPRESSION memory usage scales proportionally to sample size as well with lower fixed overhead.

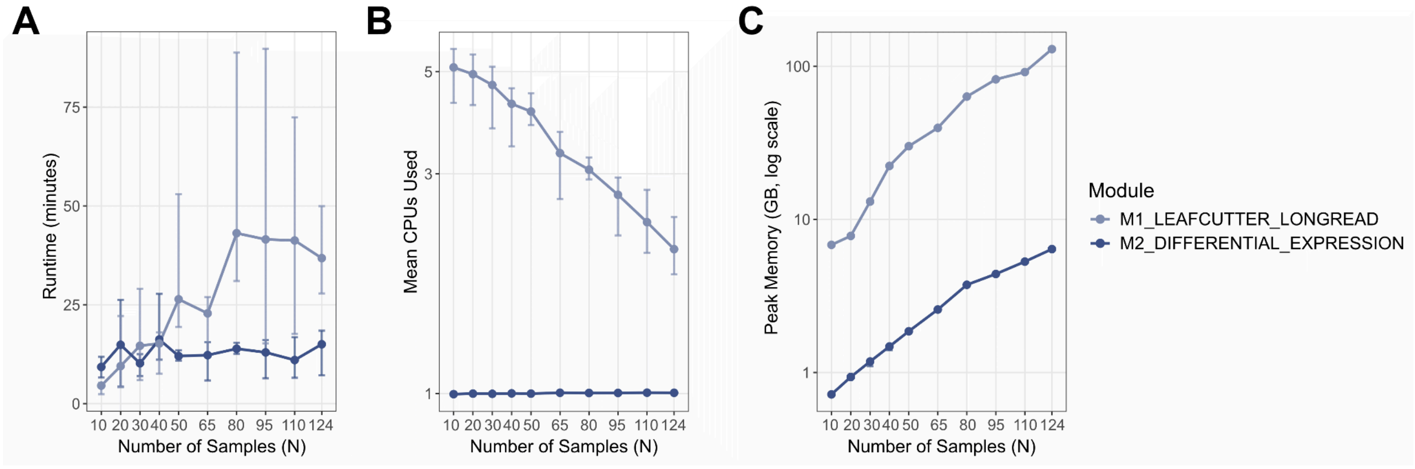


**Figure N1.6.** Subworkflow S4 Multisample Analysis.

### Note N2: Long-read adaptation of LeafCutter

LRP2 Subworkflow S4 includes a preliminary implementation of our long-read adaptation of LeafCutter (Li et al. 2017). In the original LeafCutter, the first step clusters overlapping introns using junction-spanning short reads (Figure N2.1). Each junction in a cluster has a read count. For quantification, junction usage is calculated as the junction read count divided by the cluster read count (i.e., the sum of the junction read counts). For differential usage analysis between conditions, a Dirichlet-multinomial model is applied to junction counts per cluster.

Intron clusters are a useful representation since they can capture complex splicing (i.e., a skipped exon event that overlaps two alternative 3' splice sites as in cluster 1 below). However, the combinations of junctions actually present at the transcript level cannot be directly resolved.

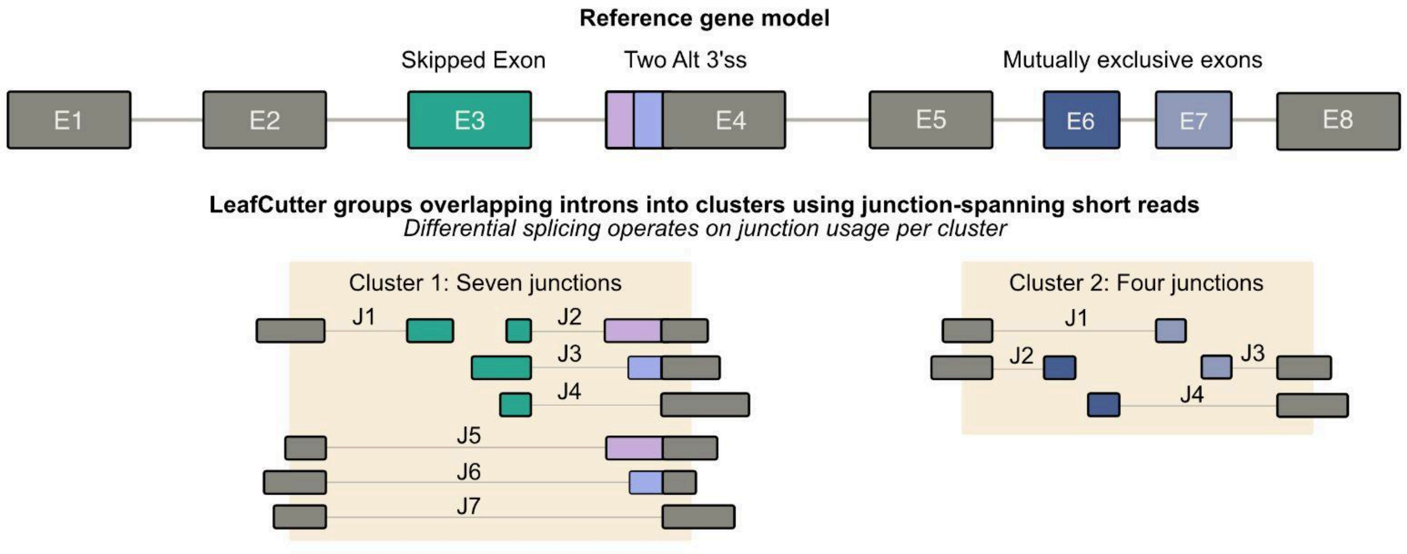


**Figure N2.1: LeafCutter applied to short read RNA-sequencing.**

Long read RNA-sequencing captures full-length transcript structures, showing which combinations of junctions occur within the same molecule. Long-read LeafCutter applies the same intron clustering as the original, then uses the long read structures to define each unique junction chain within a cluster (Figure N2.2). We call this resolution a “sub-isoform”, since it sits between the junction and the full isoform. For quantification, long read counts are collapsed to sub-isoform counts, and sub-isoform usage is calculated as the sub-isoform read count divided by the total for that cluster. For differential usage analysis between conditions, the same Dirichlet-multinomial model is applied to sub-isoform counts per cluster. By using sub-isoforms instead of junctions per cluster, we retain the broader context that long read sequencing provides, while still collapsing the complexity of full length transcripts.

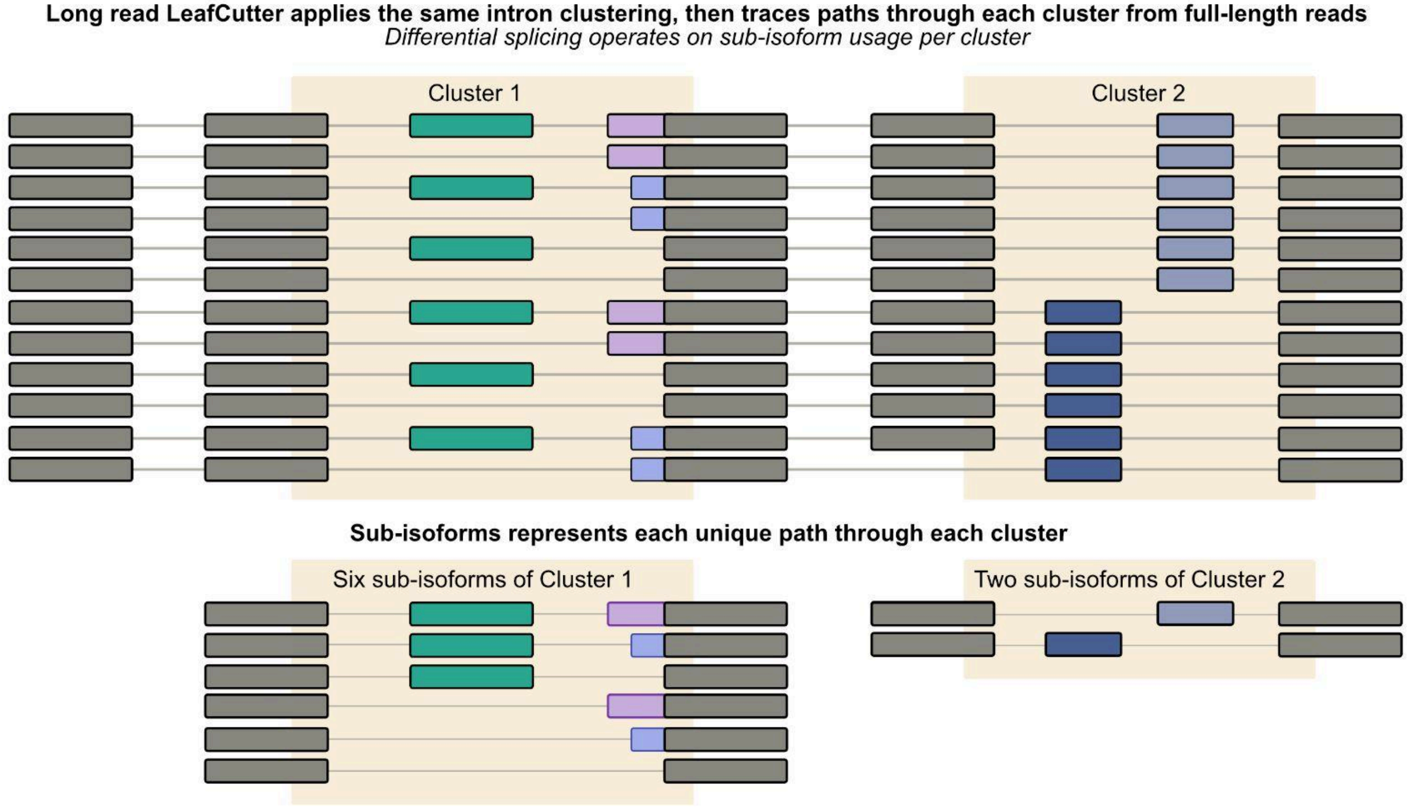


**Figure N2.2. Long read LeafCutter applied to long read RNA-sequencing**

**Note N3: Evaluation of CPAT ORF calling and LRP2 selection accuracy against GENCODE annotation**

LRP2 Subworkflow S3 uses CPAT (Wang et al. 2013) to call open reading frames (ORFs) for the refined transcript set output from Subworkflow S2. LRP2 runs CPAT with `--min-orf=75` and `--top-orf=5`, matching CPAT's own defaults. Comparison between CPAT and other ORF callers can be found in our original LRP manuscript and in another recent manuscript (Miller et al. 2022; Xu et al. 2026).

To evaluate CPAT's performance, we called ORFs on GENCODE v49 protein-coding transcripts and compared them to their respective annotated ORF. For 97.7% of MANE and 94.6% non-MANE transcripts, CPAT reported the annotated ORF (unfiltered, considering all 5 ORFs/transcript; Figure N3.1). Out of the 2.3% and 5.4% that CPAT missed, the majority were cases where there is an upstream ATG that shares the same stop codon as the annotated ORF. By design, CPAT misses these because it only reports the longest ORF per stop codon. Importantly, the coding region and frame are correct, but there is an N-terminal extension. CPAT also only reports ATG starts. Finally, as described in (Miller et al. 2022), many of the "missed:other" transcripts are selenocysteine recoding events, in which the stop codon is recoded to selenocysteine.

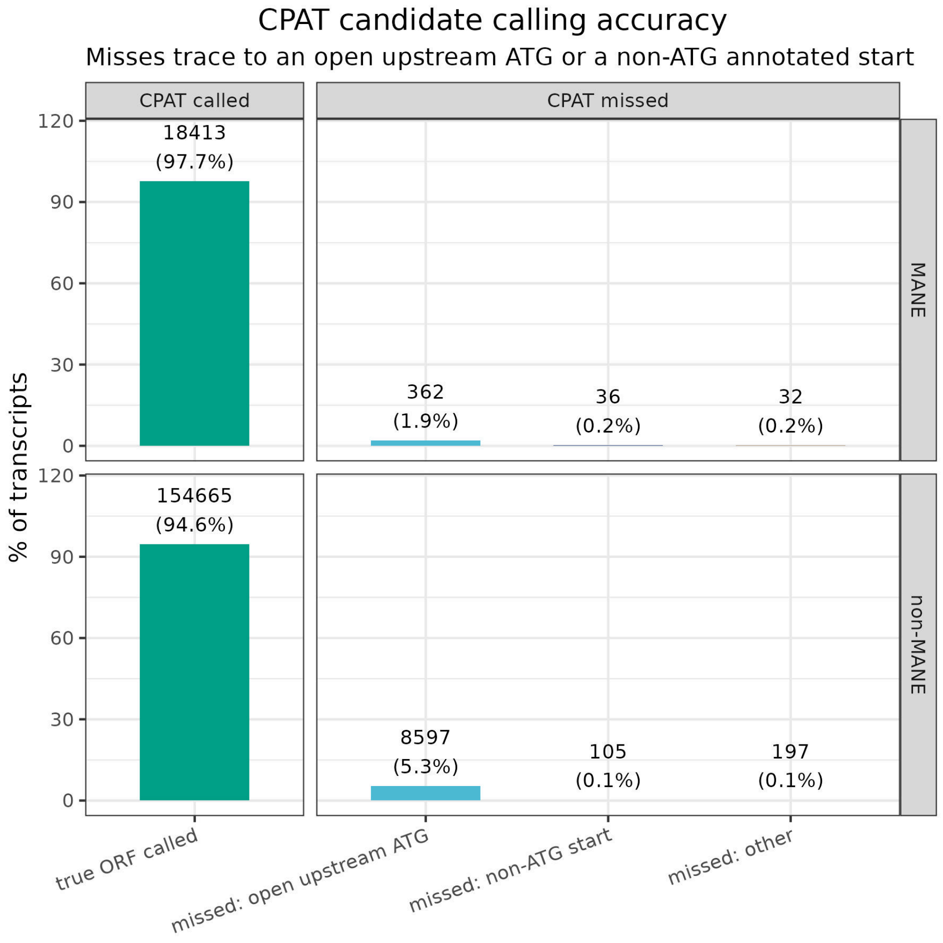


**Figure N3.1**

From all of the top candidate ORFs that CPAT calls, LRP2 selects the best-supported ORF per transcript, using criteria outlined in the original LRP pipeline (Miller et al. 2022).

1. An ORF must first pass CPATs coding probability threshold (plausible ORF > 0.364 for humans)
2. If a plausible ORF matches a GENCODE annotated ORF, that ORF is selected (this criteria is bypassed for this evaluation)
3. All other plausible ORFs must have a stop codon
4. For the remaining plausible ORFs, the ORF whose start codon is furthest upstream is selected based on the ribosomal scanning model of translation (Kozak 1999)

For the GENCODE v49 transcripts, LRP2 criteria selected the correct ORF as the best supported ORF for 94.7% and 90.5% of MANE and non-MANE transcripts, respectively (Figure N3.2). In the cases where LRP2 did not select the annotated ORF, 2.3% and 5.4% are due to CPAT missing the ORF as discussed above (Figure N3.1). The remaining transcripts were split between the coding probability filter removing the true ORF (1.9% and 2.3%) and the upstream ATG ranking selecting an alternative ORF (1.2% and 1.7%).

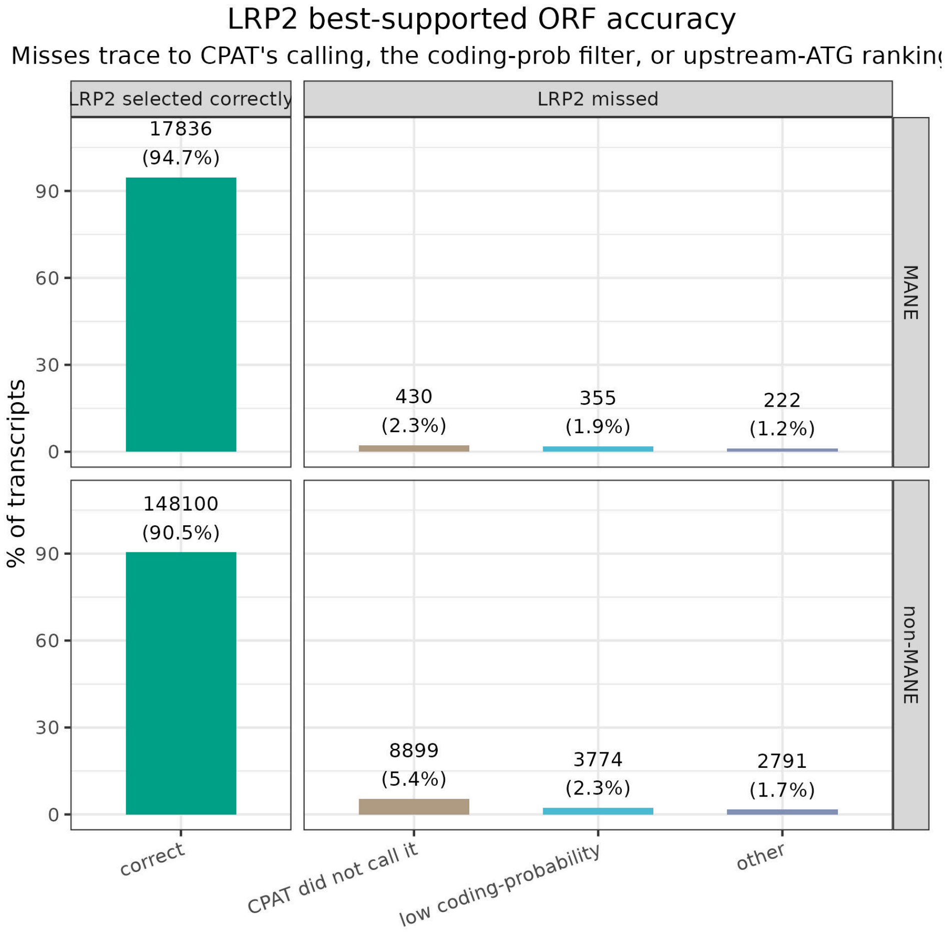


**Figure N3.2**

LRP2 applies the CPAT recommended coding probability threshold (> 0.364 in humans). We evaluated selection accuracy at different thresholds and found that between thresholds of 0.2 to 0.5, accuracy is stable in both MANE and non-MANE sets (accuracy varies by <1 percentage point). However, lowering or removing this filter altogether allows low-scoring ORFs to outrank the correct ORF in our GENCODE annotated ground truth set (Figure N3.3).

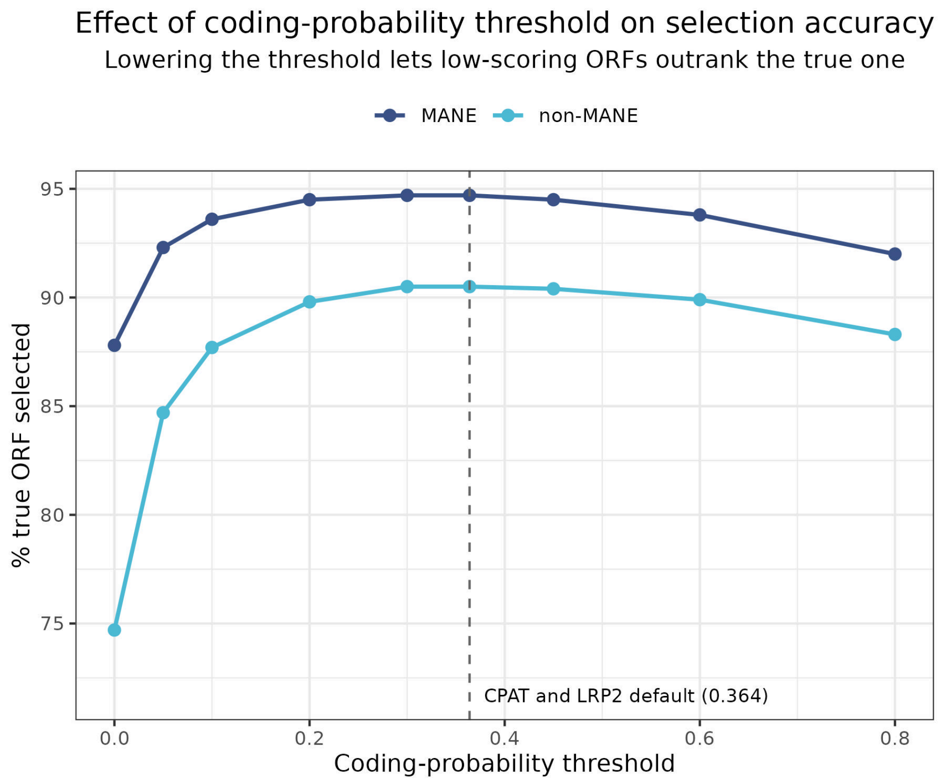


**Figure N3.3**
